# Liver humanized NSG-PiZ mice support the study of chronic hepatitis B virus infection and antiviral therapies

**DOI:** 10.1101/2022.05.17.492375

**Authors:** Rossana Colón-Thillet, Daniel Stone, Michelle A. Loprieno, Lindsay Klouser, Pavitra Roychoudhury, Tracy K. Santo, Hong Xie, Larry Stensland, Sarah L. Upham, Greg Pepper, Meei-Li Huang, Martine Aubert, Keith R. Jerome

**Affiliations:** Vaccine and Infectious Disease Division, Fred Hutchinson Cancer Center, Seattle, WA, USA; Department of Laboratory Medicine and Pathology, University of Washington, Seattle, WA, USA

**Keywords:** Hepatitis B virus, engraftment, cccDNA, hepatocyte, adeno-associated virus, RTi

## Abstract

Hepatitis B virus (HBV) is a pathogen of major public health importance that is largely incurable once a chronic hepatitis B (CHB) infection is established. Only humans and great apes are fully permissive to HBV replication, and this species restriction has impacted HBV research by limiting the utility of small animal models of HBV. To combat the species restriction of HBV and enable more HBV studies *in vivo*, liver-humanized mouse models have been developed that harbor primary human hepatocytes (PHH) and are fully permissive to HBV infection and replication. Unfortunately, these models can be difficult to establish and are expensive commercially, which has limited their academic use. As an alternative mouse model to study HBV, we evaluated liver-humanized NSG-PiZ mice and showed that they are fully permissive to HBV and can develop CHB. Mice were infected with a precore mutant clinical isolate that has now been serially passaged through 3 generations of mice without loss of fitness. HBV selectively replicates in hCK18+ human hepatocytes within chimeric livers, and HBV+ mice secrete infectious virions and HBsAg into blood, while also harboring covalently closed circular DNA (cccDNA). HBV+ mice remain viremic for at least 169 days, which should enable the study of new curative therapies targeting CHB and respond to antiviral entecavir therapy. The extended duration of viremia is sufficient to enable the study of established and new therapeutic approaches targeting CHB. Furthermore, HBV+ PHH in NSG-PiZ mice can be transduced by the hepatotropic AAV3b and AAV.LK03 vector capsids, which should enable the study of curative gene therapies that target CHB. In summary, our data demonstrates that liver humanized NSG-PiZ mice can be used as a robust and cost-effective alternative to existing CHB models and may enable more academic research labs to study HBV disease pathogenesis and antiviral therapy in a setting that is fully permissive to ongoing replication.

## Introduction

Hepatitis B virus (HBV) remains a major cause of mortality and morbidity globally, even though global vaccine coverage with three doses of one of several highly effective vaccines is estimated to be 83% ^1^. The WHO estimates that only 10% of people infected with HBV are aware of their infection, and that only 22% of these people are receiving antiviral therapies such as reverse transcriptase inhibitors (RTi) that could potentially limit transmission ^2^. This has led to continued spread of the disease, particularly amongst children where vaccination rates are closer to 43%, and rates of mother-to-child transmission remain higher than desired ^3^. It is estimated that 1.5 million *de novo* HBV infections still occur annually, and since chronic hepatitis B (CHB) is a lifelong condition, it is likely that close to 300 million people currently have CHB ^2–4^. This high global incidence causes approximately 820,000 HBV-related deaths each year ^2^, as patients with advanced CHB are more likely to develop complications that may lead to death, such as cirrhosis or hepatocellular carcinoma (HCC). Notably, HBV is the second leading cancer-causing virus after human papilloma virus (HPV), and it is estimated that 3% of annual global cancer deaths are directly attributable to HBV ^5^. Thus, HBV remains a major public health concern and a cure is desperately needed.

The study of HBV, both *in vitro* and *in vivo,* has been confounded by species restriction for HBV replication, since only hepatocytes from humans, great apes and Asian tree shrews (*tupaia belangeri*) are fully susceptible to HBV infection ^6^. The differences in susceptibility across species are largely (although not entirely) driven by differences in the sodium taurocholate co-transporting polypeptide (NTCP), the cellular receptor for HBV ^7–13^, which directly interacts with the Pre-S1 protein of HBV virions to mediate viral attachment and uptake. Consequently, there are significant limitations to what can be learned about HBV disease pathogenesis and treatment in many of the available cell culture and animal models of HBV infection. Since chimpanzee research has been phased out over the last 10 years, there are no animal models available to study HBV that truly replicate the clinical situation in humans where continuous virus replication and spread into uninfected hepatocytes occur in the presence of a functional host immune system. Nevertheless, a wide range of different avian, mouse, woodchuck and non-human primate (NHP) models have been developed that recapitulate different aspects of the HBV life cycle have been developed, and they are being used widely to study HBV replication and therapy (Reviewed in ^6, 14–17^).

In recent years, immune deficient liver humanized mouse models were developed that have become the gold standard for study of actively replicating HBV and antiviral therapies that target replication and spread of HBV *in vivo*. Livers in these mice are fully permissive to HBV replication and they can harbor chronic HBV infections that last the lifetime of a mouse, enabling their use to monitor the effects of novel antiviral drugs. Mice with genetic backgrounds that selectively kill mouse hepatocytes, including TK-NOG ^18^, uPA-RAG ^19^, uPA-SCID/PXB, ^20, 21^ and FRG ^22^ mice, are used to create an environment that is receptive to primary human hepatocyte (PHH) engraftment following transplantation. In these mouse models, PHH engraftment levels are sufficient to support high level replication of both laboratory and clinical HBV isolates ^19, 23–27^, enabling their use in the study of antiviral therapies ^25, 28–32^. Unfortunately, these models are technically difficult to establish, and this has limited how widely they are used for research. Ultimately, simpler and more cost-effective liver humanized mouse models are desperately needed to enable more widespread assessment of new translational therapies targeting HBV.

Studies with liver humanized mice can be prohibitive to many academic research labs since it is expensive to buy animals that have been humanized commercially, and establishment of a breeding colony in house can be difficult and may require a breeding license. For example, TK-NOG and uPA transgenic mice are challenging to breed due to male infertility or neonatal fatality due to constitutive uPA expression, whereas FRG mice require a breeding license and extensive cycled treatment with NTBC plus pretreatment with a uPA-expressing adenovirus vector to stimulate high level engraftment ^6, 15^. Therefore, in continuation of our work developing antiviral gene editing therapies targeting HBV ^31, 33^, we investigated the recently described liver-humanized NSG-PiZ mouse as an alternative model to study HBV disease pathogenesis and therapy. NSG-PiZ mice can be bred easily, need no excessive animal husbandry, and only require injection with a hepatotoxic agent to precondition the liver prior to intrasplenic PHH transplantation ^34, 35^.

Here we show that humanized NSG-PiZ mice are fully permissive to HBV replication. HBV+ NSG-PiZ mice can be followed longitudinally for at least 169 days and can be monitored for levels of human albumin (huAlb), HBsAg and viral loads, or for intra-hepatic viral DNA levels at necropsy. HBV+ mice are responsive to traditional RTi therapy, and HBV+ PHH can be transduced by hepatotropic AAV vector capsids, which is of particular importance for the study of novel gene therapies that target HBV ^36, 37^. When taken together, our data demonstrates that liver humanized NSG-PiZ mice are a robust and comparatively inexpensive model that can be readily used to study the pathogenesis of CHB and antiviral therapies targeting ongoing CHB infections.

## Materials and Methods

### Cell culture

Human embryonic kidney (HEK) 293 cells ^38^ were grown in DMEM (Thermo Fisher, Waltham, MA) supplemented with 10% fetal calf serum.

### AAV vectors

AAV vectors were generated by transiently transfecting HEK293 cells using PEI according to the method of Choi et al ^39^. Briefly, HEK293 cells were transfected with AAV vector plasmid pscAAV-smCBA-GFP ^40^, in combination with the plasmid pRepCap3b which contains Rep and Cap from AAV serotype 3b (Kindly provided by Dr. David Russell ^41^) or a plasmid that contains the AAV2 rep and AAV-LK03 capsid proteins, and a helper plasmid that expresses adenovirus helper proteins. At 24 hours post-transfection media was changed to serum-free DMEM, and after 72 hours cells were collected and re-suspended in AAV lysis buffer (50mM Tris, 150 mM NaCl, pH 8.5) before freeze-thawing 4 times. AAV stocks were purified by iodixanol gradient separation ^39, 42^ followed by ultrafiltration and concentration into PBS using an Amicon Ultra-15 column (EMD Millipore, Burlington, MA) before storage at -80°C. All AAV vector stocks were quantified by quantitative PCR using primers against the AAV inverted terminal repeat, with linearized plasmid DNA as a standard, according to the method of Aurnhammer et al^43^. AAV stocks were treated with DNase I (Thermo Fisher) and Proteinase K (Thermo Fisher) prior to quantification.

### Animal welfare statement

All animals were housed at the Fred Hutchinson Cancer Center and all experimental procedures performed were reviewed and approved by the Institutional Animal Care and Use Committee of the Fred Hutchinson Cancer Center (protocol no. 51064). This study was carried out in accordance with the recommendations in the Guide for the Care and Use of Laboratory Animals of the National Institutes of Health (“The Guide”).

### Mice

Male NSG-PiZ [NOD.Cg-*Prkdc^scid^ Il2rg^tm1Wjl^* Tg(SERPINA1*E342K)#Slcw/SzJ] mice were obtained from Jackson Laboratories (Bar Harbor, ME; strain # 028842). Mice were housed in a ABSL2 facility and received standard housing, diet, bedding, enrichment and light/dark cycles. Mice weights were monitored for the duration of each experiment.

### Liver humanization

Livers of NSG-PiZ mice were humanized according to the method of Borel *et al* with modifications ^35^. Briefly, mice were pretreated to create an environment amenable to human hepatocyte engraftment via IP injection with 100μl of the hepatotoxin monocrotaline (50 mg/kg) at 7 and 14 days prior to human hepatocyte transplant, or via retro-orbital injection with 1μg of an anti-CD95 antibody on the day of human hepatocyte transplant. Mice were administered Buprenorphine SR on the day prior to surgery to provide ∼72 hours of analgesia. On the day of transplant, 6–10-week-old mice were anaesthetized with isoflurane then an incision was made in the left flank below the rib cage to expose the spleen. A total of 5×10^5^ primary human hepatocytes were then slowly injected into the exposed spleen in a volume of 50μl of sterile 1X Hanks Balanced Saline (ThermoFisher, Waltham, MA,) using a 27G 0.5ml insulin syringe (Easy Touch, Houston, TX). The needle was left in the spleen for 30 seconds after injection before slow removal, and pressure was applied to the injection site with a sterile cotton swab to limit bleeding. The spleen was then returned to the abdominal cavity which was closed with a 4-0 Vicryl suture (Ethicon, Cincinnati, OH), and the skin closed with wound clips.

### Human hepatocytes

Primary human hepatocytes were obtained from BioIVT (Westbury, NY) from a 10-month-old male, African American donor (donor A) and a 3-year-old, male Caucasian donor (donor B). Cryo-preserved cells were thawed according to the supplier’s instructions before resuspension in sterile 1X Hanks Balanced Saline.

### HBV isolate

HBV+ serum was obtained from BioIVT (Westbury, NY; human serum/HMN13090). To determine the genotype and consensus sequence of the clinical isolate used in our study we used a probe capture approach previously described for capture of HSV and HHV-6 ^44, 45^ to obtain a >10X coverage consensus HBV sequence. Briefly, tiled probes (XGen Custom Hybridization Probe Panel, IDT, Coralville, IA) were generated against the NCBI reference HBV genome (Genbank Accession NC_003977) and used to capture HBV DNA which was then sequenced via NGS. Significant heterogeneity was seen at multiple locations within the HMN13090 HBV genome suggesting a diverse viral quasispecies was present within our ‘founder’ clinical isolate. HMN13090 has been serially passaged through 3 generations of NSG-PiZ mice, using serum pooled from HBV+ mice to create a challenge inoculum for subsequent mice that was tittered by qPCR as indicated below. For HBV challenge, HBV+ serum diluted in sterile 1X PBS was administered to at a dose of 1 x 10^6^ genome equivalents (copies) per animal via retro-orbital injection in a volume of 50μl. The sequence for HBV isolate HMN13090 has been uploaded to Genbank (Accession # OM194175) and contains an in-frame TGG to TAG (W to STOP) mutation in PreCore that is highly prevalent in genome sequences from the HBVdb database ^46^. The W28* mutation stabilizes the HBV stem loop ^47^ and was previously implicated in the development of cirrhosis and HCC in advanced disease^48^. M2 mutations have been shown to emerge shortly before or around the time of HBe seroconversion^47, 49, 50^.

### AAV gene transfer

scAAV.LK03-smcBA-GFP or scAAV3B-smCBA-GFP vectors were delivered to naïve (not humanized) NSG-PiZ mice at 10^12^ vector genomes per mouse (n=1) or to HBV-positive liver humanized NSG-PiZ mice at doses of 2 x 10^10^, 2 ×10^11^ or 1 x 10^12^ vector genomes per mouse (n=3/dose) via retro-orbital injection in a volume of 50μl diluted in USP 1X PBS. Mice were euthanized 28 days later to analyze liver transduction.

### Drugs and antibodies

Monocrotaline was obtained from Oakwood Chemicals (Estill, SC; item # 002602) and 187.5 mg was dissolved in 1.8 ml of 1 M HCl followed by addition of 3 to 4 ml of distilled water. This solution was adjusted to pH 7.4 using 1 M NaOH solution and filled up to 15 ml with distilled water to make a stock solution of 12.5 mg/mL. Purified NA/LE hamster anti-mouse anti-CD95 antibody was obtained from BD Biosciences (clone Jo2 (RUO); catalogue # 554254). Entecavir was obtained from Toronto Chemicals (Toronto, Ontario, Canada; catalogue # E558900), was resuspended in Medidrop sucralose (Clear H2O; product code # 75-01-1001) and administered orally at 0.5 mg/Kg/day. Ciclopirox was obtained from Bio-Techne/Tocris (Minneapolis, MN; Catalogue # 6384), was resuspended in 4% ethanol, 5.2% Tween 80 and 5.2% PEG-400 in PBS and administered daily at 5 mg/kg via IP injection.

### Human albumin quantification

Human albumin levels in mouse sera were determined using the Human Albumin ELISA Kit (Bethyl Laboratories, E88-129).

### HBsAg and HBeAg quantification

HBsAg quantifications were performed on the Abbott Architect i2000 using the HBsAg qualitative (Abbott Laboratories, Des Plaines, IL; product #B4P530) and HBsAg qualitative confirmatory (Abbott Laboratories, Des Plaines, IL; product #B4P540) systems. Anti-HBeAg antibodies and HBeAg were quantified using the ETI-AB-EBK PLUS (anti-HBe) kit (Diasorin, Cypress, CA;product # P001929) or the ETI-EBK PLUS (HBeAg) kit (Diasorin, Cypress, CA;product # P001930) respectively.

### Quantitative PCR

Total HBV DNA was detected in DNA extracted from serum using the MagNA Pure 96 system (Roche, Basel, Switzerland). Total HBV DNA, HBV cccDNA, human RPP30, and mouse RPP30 were all detected in DNA extracted from chimeric humanized NSG-PiZ mouse livers. For total HBV DNA, human RPP30, and mouse RPP30, ddPCR was performed using genomic DNA extracted using the DNeasy Blood and Tissue kit (Qiagen, Hilden, Germany). For cccDNA, ddPCR was performed using genomic DNA extracted via a modified HIRT procedure as described below, followed by treatment with T5 exonuclease (NEB, Ipswich, MA) at 37°C for 1 hour. Primer/probe sets for HBV total DNA ^31^, human RPP30 ^33^, mouse RPP30 ^31^ and cccDNA ^51^, have been described previously.

### Modified Hirt extraction

The modified HIRT extraction was performed using a previously described protocol ^52^. Briefly, homogenized liver tissue extracts underwent a neutral pH, 3 step alkaline lysis before the supernatant was passed over a Qiagen mini prep column to capture extra-chromosomal DNA.

### Liver chimerism

The levels of hepatocyte chimerism (the percentage of human hepatocytes within all hepatocytes) were determined for individual mice by determining the levels of human and mouse RPP30 in humanized livers by ddPCR, then extrapolating the percentage of human hepatocytes as previously described ^31^. This method assumes that all human RPP30 signal is derived from PHH and that the mouse RPP30 signal is derived from murine hepatocytes, Kupffer cells, sinusoidal endothelial cells, hepatic and stellate cells at levels previously described for liver humanized uPA-SCID mice ^53^.

### Histopathology and Oil Red O staining

Mice were perfused with 1X PBS at necropsy and tissues were either snap frozen for DNA/RNA extraction, placed in 4% PFA overnight prior to embedding in paraffin or post fixed in 4% PFA at 4°C overnight before then cryopreserving in 30% sucrose at 4°C overnight and embedding in OCT. Paraffin-embedded tissues were sectioned at 5μm and then stained with hematoxylin and eosin. Frozen tissues were sectioned at 8μm and stained with Oil Red O by the Fred Hutchinson Cancer Center, Experimental Histopathology Core. Oil Red O is a fat-soluble dye that stains neutral triglycerides and lipids in frozen sections, and is routinely used to detect steatosis in liver sections ^54^.

### Immunohistochemistry

Immunohistochemistry (IHC) was performed on 5μm paraffin sections. Sections were progressively hydrated in xylene, 100% ethanol, 95% ethanol, 70% ethanol and 50% ethanol, then treated with citrate antigen retrieval buffer (10 mM Sodium citrate, 0.05% Tween 20, pH 6.0) at 95°C for 30 minutes and blocked with 10% donkey serum with 1% BSA for 1 hour. For staining of HBV surface antigen (HBsAg), human cytokeratin 18 (hCK18) and GFP, sections were incubated with rabbit polyclonal anti-HBsAg (Bio-Rad, Hercules, CA, catalogue # OBT0990), mouse anti-hCK18 (Agilent, Santa Clara, CA, DC10 catalogue # M701029-2) and goat polyclonal anti-GFP (Novus Biologicals, Centennial, CO, catalogue # NB100-1770). All primary antibody incubations were done in in TBS with 1% BSA at 4°C overnight. Sections were washed 2 times with TBS + 0.025% triton then incubated with secondary antibodies donkey anti-mouse Alexa Fluor 647 (ThermoFisher, Waltham, MA, catalogue # A31571), donkey anti-rabbit Alexa Fluor 594 (ThermoFisher, Waltham, MA, catalogue # A21207) and anti-goat Alexa Fluor 488 (Abcam, Cambridge, UK catalogue #ab150129) at room temperature for 1 hour. Sections were counterstained with DAPI before mounting with Prolong Gold Antifade Reagent (Thermo Fisher, Waltham, MA catalogue #P36934).

### Microscopy

Photomicrographs were obtained from the Halolink platform (Indica Labs) using images acquired at 20X or 40X using the Leica Biosystems Aperio VERSA 200 slide scanner.

## Results

### Kinetics of PHH engraftment and HBV infection in NSG-PiZ mice

To determine whether liver humanized NSG-PiZ mice can support the study of chronic hepatitis B infection we simultaneously determined the long-term kinetics of primary human hepatocyte (PHH) repopulation and HBV replication within the livers of HBV-infected and naïve NSG-PiZ mice (**Figure 1A**). A total of 25 mice were pre-conditioned with the hepatotoxic plant-derived pyrrolizidine alkaloid monocrotaline (MCT) then transplanted with PHH from two different pediatric donors to account for potential donor variability in engraftment capacity (Donor A, n = 13; Donor B, n = 12). As a surrogate for liver repopulation by PHH, we assessed human albumin (huAlb) levels in mouse sera at 8 weeks post-engraftment for all transplanted mice. In the donor A group, 12 out of 13 mice had detectable levels of huAlb ranging from 0.002 to 2.34mg/mL, whereas in the PTC group 12 out of 12 mice had detectable huAlb levels ranging from 0.12 to 4.11mg/mL (**Figure 1B**).

**Figure 1.**
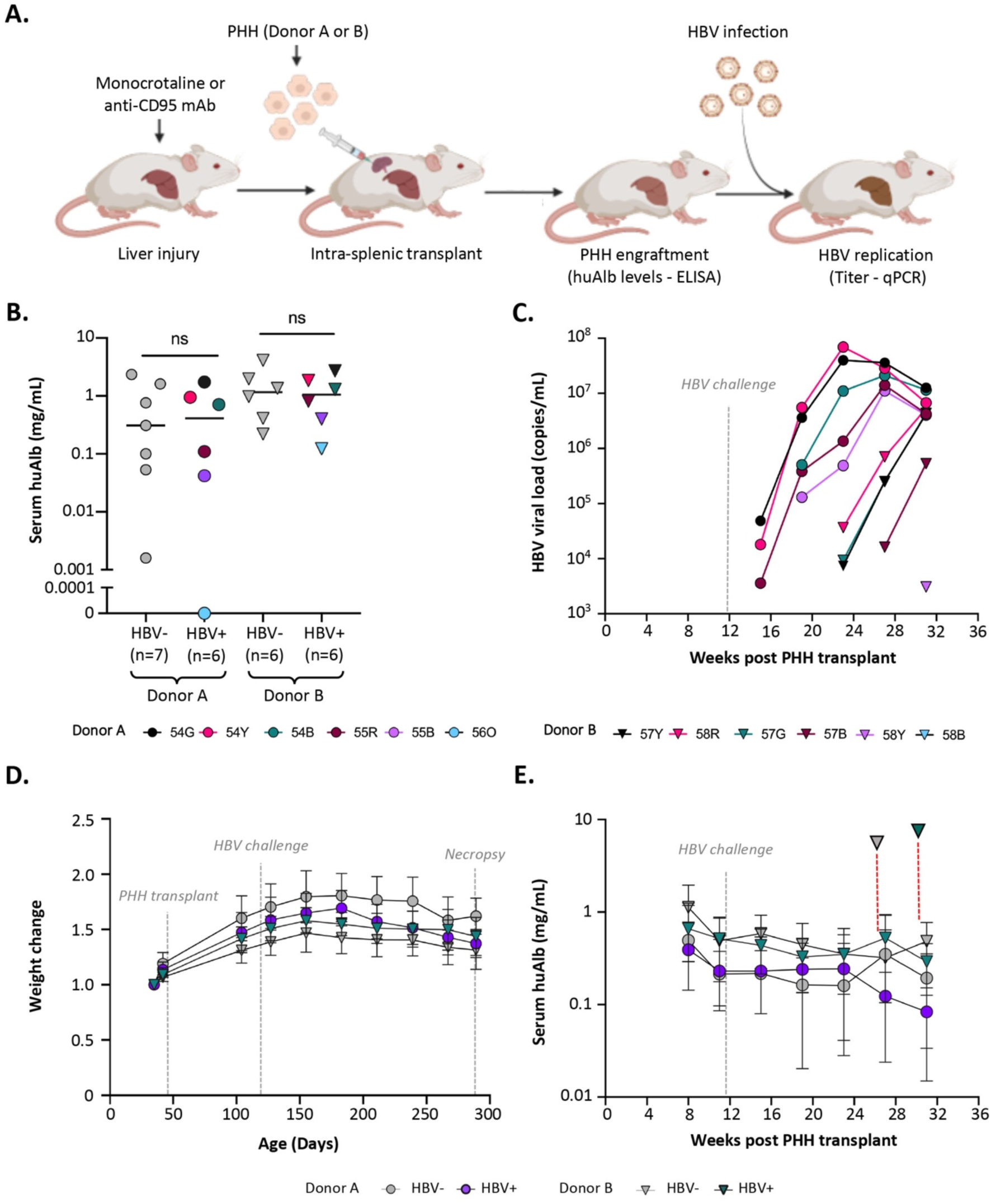
Longitudinal pilot study of PHH engraftment and HBV replication in liver humanized NSG-PiZ mice. (**A**) experimental scheme. A total of 25 mice were transplanted with PHH from 2 pediatric donors for this pilot study. (**B**) Experimental grouping was based on serum huAlb levels in monocrotaline pretreated and PHH transplanted mice at 8 weeks post-transplant. huAlb levels were equalized by donor for HBV negative (grey symbols) and positive (colored symbols) mice prior to challenge with HBV. (**C**) HBV viral loads in serum following intravenous challenge with 10^6^ infectious units of a genotype D clinical isolate. HBV challenged mice are color coordinated between panels B and C. (**D**) Weight change over time. (**E**) Longitudinal huAlb levels. One mouse from each group transplanted with donor B PHH died prematurely at days 183 and 215 post-transplant and both are indicated in panel E (dotted red lines).

Next, we asked whether PHH grafts remain stable in livers of HBV-infected and HBV-uninfected NSG-PiZ mice. We used huAlb levels to distribute mice into four experimental groups (Donor A-uninfected, Donor A-infected, Donor B-uninfected, Donor B-infected) to normalize for the variability in the degree of engraftment by donor (**Figure 1B**) and intravenously challenged mice in groups Donor A-infected and Donor B-infected with a genotype D HBV clinical isolate at 11 weeks post PHH-transplant. Sera from all groups was collected every four weeks and analyzed for levels of huAlb and HBV DNA. HuAlb levels peaked at 8 weeks post-transplant in all groups, then slowly declined over time as expected (**Figure 1E**) given previous work showing that Z-AAT levels in NSG-PiZ decrease as mice age and the selective growth advantage this provides PHH is reduced over time ^35^. Mice receiving PHH from donor B (infected and uninfected) had slightly higher levels of huAlb than mice receiving PHH from donor A (∼2- fold on average) at all timepoints. In groups Donor A-infected and Donor B-infected 5 of 6 mice became viremic. Viral loads were highest (up to 7×10^7^ copies/mL) in mice with the highest huAlb levels at 8 weeks post PHH transplant (**Figure 1C**). HBV DNA was detected earlier in sera from mice with higher huAlb levels for both PHH donors and was not detected at any time point in mice with huAlb levels below 0.04mg/mL at 8 weeks post PHH transplant. Unexpectedly, the onset of viremia was delayed by 4-8 weeks and peak viral loads were lower for mice from group donor B-infected, despite having higher engraftment levels than mice from group donor A-infected (**Figure 1C, Figure S1**). No significant difference in huAlb levels was seen between infected and uninfected mice over time, and for both PHH donors, the time from HBV detection to peak viremia was 8-12 weeks. For the duration of the study, mouse weights remained stable in all treatment groups (**Figure 1D**).

### Histological analysis of chimeric mouse livers

We analyzed the left lateral, right medial and left medial lobes for H&E stained livers from PHH-naïve Swiss-Webster (PHH-), PHH-naïve NSG-PiZ (PHH-), anti-CD95 antibody treated PHH-naive NSG-PiZ (PHH-), MCT treated PHH-naïve NSG-PiZ (PHH-), humanized NSG-PiZ (PHH+/HBV-), and HBV-infected humanized NSG-PiZ (PHH+/HBV+) mice for gross pathology and histopathology. Livers from PHH-naïve NSG-PiZ mice and PHH-naïve NSG-PiZ mice pre-treated with anti-CD95 antibody or MCT showed abnormal hepatocyte pathology that was not seen in PHH-naïve Swiss-Webster mice (**Figure 2A**). There were also signs of inflammation and loss of endothelial integrity in PHH-naïve NSG-PiZ and anti-CD95 antibody or MCT pre-conditioned livers. Humanized NSG-PiZ mouse livers (PHH+/HBV-) had a distinctive appearance in each lobe after H&E staining similar to humanized uPA-SCID and TK-NOG mice ^21, 25, 26, 53^ that was independent of the pre-conditioning method or PHH donor. We identified clusters of hepatocytes with a clear cytoplasm and a granular appearance that frequently contained large holes and were interspersed between darkly stained regions of hepatocytes with normal appearance (**Figure 2B**). The clear hepatocytes appeared to have undergone microvesicular or macrovesicular steatosis, and stained positive for lipid accumulation by Oil red O (**Figure 2C**). Humanized NSG-PiZ mouse livers stained with H&E did not differ in their gross appearance when infected with HBV. Livers from PHH-/HBV-, PHH+/HBV- and PHH+/ HBV+ NSG-PiZ mice were also stained with picrosirius red to visualize collagen and determine whether fibrosis occurs in humanized NSG-PiZ mouse livers (**Figure 2D**). In PHH-/HBV- mice, collagen staining was restricted to the borders of blood vessels. In contrast, collagen deposits in both PHH+/HBV- and PHH+/ HBV+ mice were also detected in highly vascularized areas of PHH engraftment.

**Figure 2.**
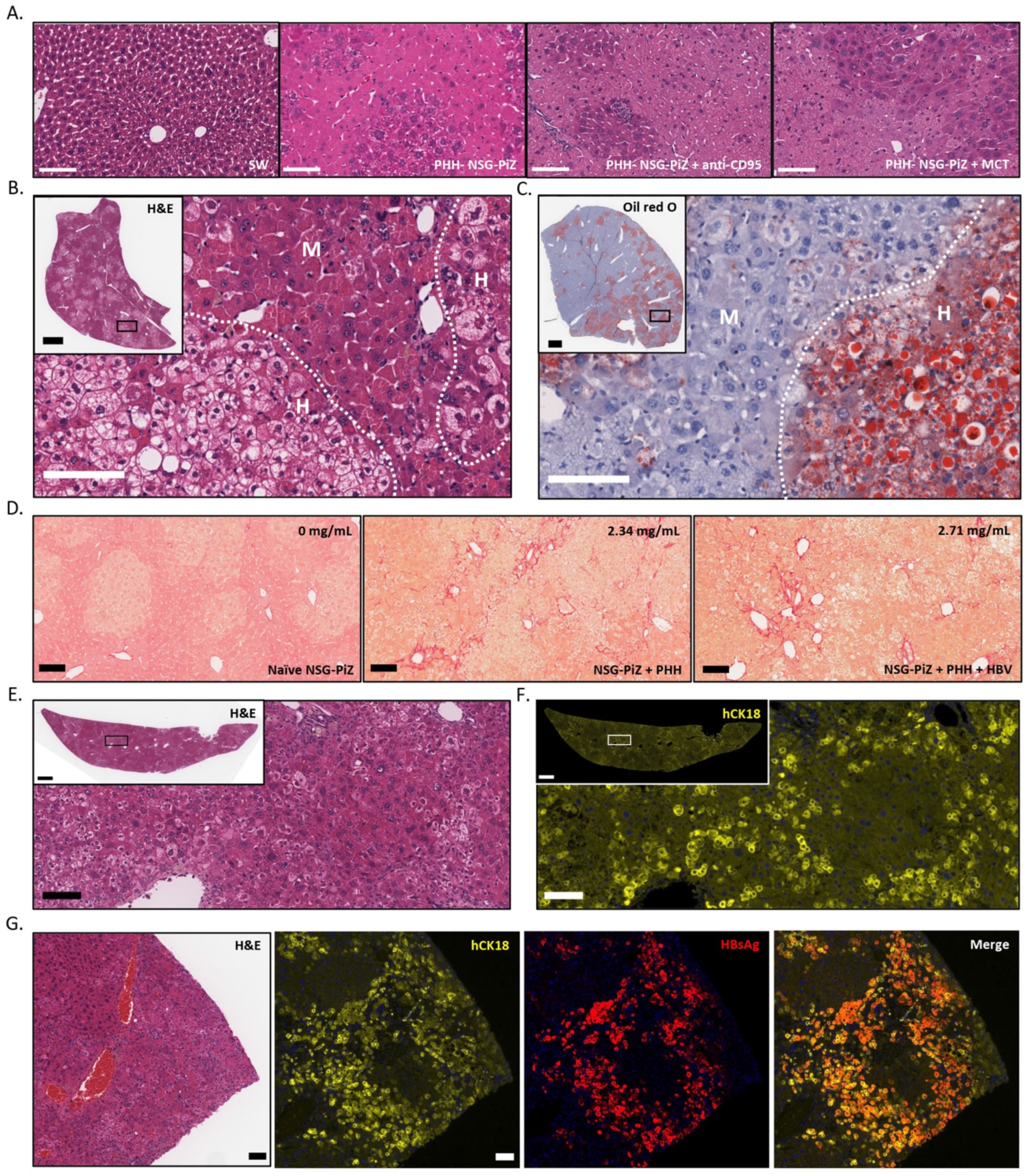
Control and humanized mouse liver phenotype. (**A**) H&E staining of control livers from a PHH- Swiss-Webster mouse, a PHH- NSG-PiZ mouse, a PHH- NSG-PiZ mouse that was euthanized 3 hours after IV delivery of 1mg of anti-CD95 mAb, or a PHH- NSG-PiZ mouse euthanized 7 days after two IP injections of monocrotaline (50mg/Kg) delivered 7 days apart). Scale bars 100μm. H&E (**B**) or Oil Red O (**C**) stained liver from an NSG-PiZ mouse pretreated with anti-CD95 mAb then transplanted with 10^6^ PHH that was euthanized at 139 days post-transplant. Approximate borders (dotted line) between human (H) and mouse (M) hepatocytes are indicated. Scale bar 100μm (main image) or 1mm (inset image). (**D**) Picrosirius red staining of livers from PHH- NSG-PiZ mice or liver humanized NSG-PiZ mice +/- HBV that had similar huAlb levels at 8 weeks post PHH transplant. HBV challenge occurred at 11 weeks post PHH transplant and humanized mice were euthanized at 242 days post-transplant (163 days post HBV challenge). Scale bars 100μm. H&E staining (**E**) and hCK18 immunofluorescence (**F**) of serial liver sections from an NSG-PiZ mouse pretreated with two IP injections of monocrotaline then transplanted with 5×10^5^ PHH that was euthanized at 233 days post-transplant. Scale bar 100mm (upper image) or 1mm (lower image). (**G**) NSG-PiZ mouse pretreated with monocrotaline, transplanted with 5×10^5^ PHHs, and challenged intravenously with 10^6^ IU of a genotype D HBV clinical isolate. HBV challenge occurred at 10 weeks post hepatocyte transplant and liver from a mouse that died at 150 days post HBV challenge is shown. Serial sections were stained with H&E or via immunofluorescence with antibodies against hCK18 (yellow) and HBsAg (Red). Scale bar - 1mm (left hand images) or 100mm (right hand images). SW – Swiss-Webster.

We next performed immunohistochemistry (IHC) on chimeric livers to detect human cytokeratin 18 (hCK18) and HBV surface antigen (HBsAg), in order to determine which hepatocytes were of human origin and whether HBV replication is restricted to PHH. The clear hepatocytes visible by H&E staining were mostly hCK18 positive (**Figure 2E/F**) except for some within the center of densely packed clusters that looked morphologically identical but were weakly positive or negative for hCK18 (**Figure S2**). In mice with lower or moderate levels of engraftment, hCK18 staining was predominantly periportal or surrounding hepatic lobules containing mouse hepatocytes, and no hCK18 positive cells were detected around the central vein (**Figure S2**). Co-labelling with HBsAg confirmed that HBV replication is restricted to hCK18-positive PHH and can be detected in all lobes of the liver (**Figure 2G, Figure S3)**, although donor-to-donor variation in the pattern of HBsAg staining was seen. In mice receiving donor A PHH, HBsAg was detected in the majority of PHH within each hCK18+ cluster, whereas in mice receiving PHH from donor B, HBsAg was only detected in distinct foci within each hCK18+ cluster, predominantly in PHH that were undergoing macrovesicular steatosis (**Figure 2F vs Figure S4**). This different pattern of HBsAg staining in donor B mice is indicative of viral spread through the cluster and could explain the delayed kinetics of viremia seen in these mice. Whether this spread of HBV into adjacent PHH is NTCP-dependent or independent is yet to be determined.

To determine how PHH donor and HBV replication impact longitudinal survival of PHH, we determined the levels of engraftment at study end for all 25 mice. Mice were euthanized at day 163 post HBV challenge (242 days post-transplant) except for two mice that died prematurely at days 106 (donor B, HBV negative) and 138 (donor B, HBV positive) post HBV challenge respectively. We scored engraftment from 0-4 using serial sections from 3 lobes of each mouse liver stained with H&E and hCK18 as shown (**Figure S5**). Although our study was not powered sufficiently to determine significance, we found that mice receiving donor B PHH had higher levels of engraftment than those receiving donor A PHH (**Figure S6**), which was expected given the ∼ 2-fold higher levels of huAlb in mice receiving donor B PHH (**Figure S1**). We also found that uninfected mice had higher levels of engraftment irrespective of the PHH donor.

### Effects of single and combination drug antiviral therapy on HBV-positive NSG-PiZ mice

We next evaluated the viral suppression effects of the nucleoside reverse transcriptase inhibitor entecavir (ETV) alone or in combination with the capsid assembly inhibitor ciclopirox (CPX) in HBV-positive liver humanized NSG-PiZ mice. CPX was included to determine whether antiviral synergy can occur when combined with ETV, as previously seen when combined with the RTi tenofovir in humanized uPA-SCID mice ^55^. For this study, we transplanted naïve NSG-PiZ mice with PHH from donors A and B after MCT or anti-CD95 pre-treatment. Levels of huAlb were determined at 12 weeks post-engraftment for all mice (**Figure 3A**). Pre-conditioning with MCT resulted in higher repopulation levels than the anti-CD95 antibody for both donors, as reported previously ^35^ (donor A-MCT vs. donor A-CD95, p = <0.0001; donor B-MCT vs. donor B-CD95, p = 0.004).

**Figure 3.**
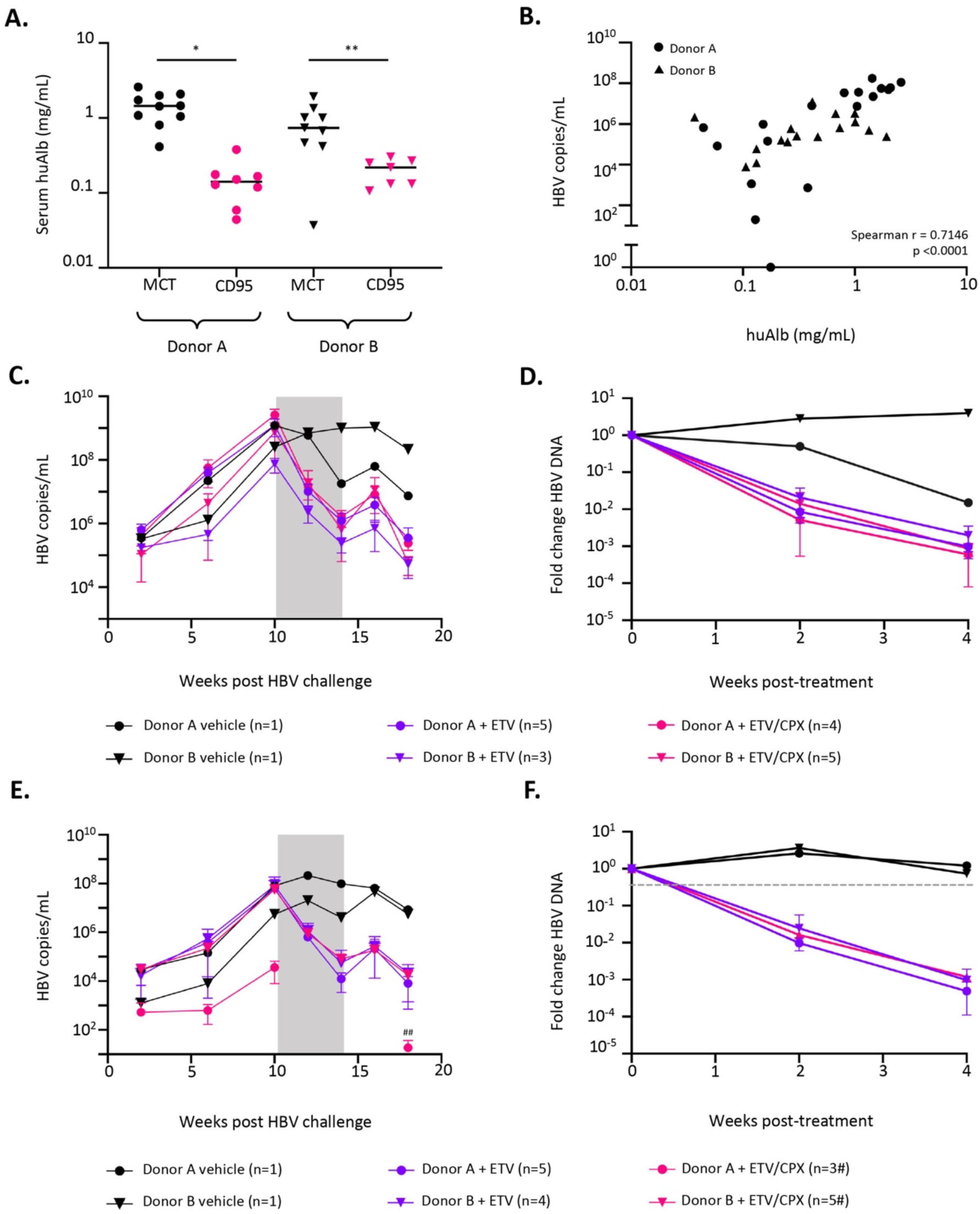
Humanized NSG-PiZ mice support HBV replication and respond to antiviral therapy. (**A**) Human albumin levels in serum were quantified via ELISA at 12 weeks after PHH transplant from Donor A or B after liver pre-injury with MCT or anti-CD95 antibodies. (**B**) Spearman correlation between human albumin levels at 12 weeks post PHH transplant and HBV viral loads at 6 weeks post-HBV infection. Longitudinal HBV viral loads in serum for mice pre-treated with MCT (**C**) or anti-CD95 antibody (**E**) were quantified by qPCR before, during, and after antiviral treatment. Fold change in HBV viral loads during treatment with ETV or ETV/CPX in mice pre-treated with MCT (**D**) or anti-CD95 antibody (**F**). At 10 weeks post-engraftment humanized NSG-PiZ mice with 10^6^ genome copies of a Genotype D clinical isolate. At 10 weeks post-infection, chronically infected mice were given ETV in drinking water (0.5mg/Kg) +/- daily IP injections of CPX (5mg/Kg) as indicated for 4 weeks (Grey bars). ^#^ 1 mouse was found dead after week 10. ^##^ virus was not detected in this group between weeks 10 and 18 post HBV challenge. *, p <0.0001; **, p = 0.004).

Mice were inoculated intravenously with HBV at 10 weeks post-engraftment and HBV titers in serum were longitudinally monitored by qPCR (**Figure 3C/E**). At 10 weeks post-infection, HBV titers reached approximately 10^8^ to 10^9^ copies/mL in the MCT group and 10^7^ copies/mL in the CD95 group. We found a strong positive correlation between the HBV titers at week 6 post-infection, the earliest time point when all mice were viremic, and huAlb levels at week 12 post-transplant (2 weeks post HBV challenge) for all mice in this experiment (n = 35), suggesting that PHH engraftment rates are positively associated with HBV titers (**Figure 3B**, Spearman r = 0.7146); this likely explains the lower levels of viremia for both donors in the anti-CD95 pretreated mice. At 10 weeks post HBV challenge mice pretreated with MCT or anti-CD95 antibody were administered ETV alone or in combination with CPX for four weeks to evaluate potential synergistic effects of HBV-targeted drug combinations on viral suppression. Treatment with ETV reduced viral titers by 1-3 log over 4 weeks for MCT and anti-CD95 pretreated mice engrafted with both donor PHH (**3D/F**). Concurrent treatment with CPX did not have a detectable synergistic effect on viral suppression (**Figure 3D/F**). After antiviral therapy cessation, HBV titers rebounded ∼1 log within two weeks but subsequently dropped another log at four weeks, although this reduction was also seen in control mice and coincided with reduced huAlb levels in humanized mice (**Figure 3C/E**).

In parallel to viral load quantification, HBeAg, anti-HBeAg antibody and HBsAg serum levels were measured in mice 10 weeks after HBV challenge, and then followed longitudinally during the antiviral treatment phase and for 2 weeks after antiviral withdrawal. Neither HBeAg nor anti-HBeAg antibodies were detected in any of the mice in this study at any time point so we analyzed the HBV genome for the presence of precore mutations by generating a consensus sequence from the challenge inoculum by NGS (Genbank accession OM194175). We found a previously-described mutation in the precore N-terminus (TTG to TAG, W28STOP) that prevents the synthesis of HBeAg and is found in 26% of 9289 HBV genomes within the HBV:Db database ^46–50^. For HBsAg, peak levels were seen at 10 weeks post HBV infection, varied across 4 orders of magnitude and were highest in MCT-pretreated mice transplanted with donor A PHH (**Figure 4A**). A strong positive correlation was seen between serum HBsAg levels and viral loads (**Figure 4B**). In all MCT pretreated and donor A transplanted mice, HBsAg levels progressively decreased at weeks 2, 4 and 6 post treatment with ETV or ETV/CPX (**Figure 4C/D**). Levels of HBsAg in 8 of 9 MCT pretreated and donor B PHH transplanted mice receiving ETV or ETV/CPX therapy briefly rose at 2 weeks post therapy then progressively decreased at weeks 4 and 6 post therapy (**Figure 4C/D**). In 7 of 8 anti-CD95 pretreated and donor A transplanted mice, HBsAg levels progressively decreased between weeks 2 and 6 post treatment with ETV or ETV/CPX (**Figure 4C/D**). In 8 of 9 anti-CD95 pretreated and donor A transplanted mice, HBsAg levels progressively decreased between weeks 2 and 6 post treatment with ETV or ETV/CPX (**Figure 4C/D**). No synergistic effect was seen in mice receiving CPX in addition to ETV irrespective of pretreatment regimen or PHH donor. Single outlier mice in 3 of 8 experimental groups made it hard to analyze group trends regarding mean HBsAg fold change over time that were clear upon comparison of individual mice (**Figure 4C-F**, **Figure S7**).

**Figure 4.**
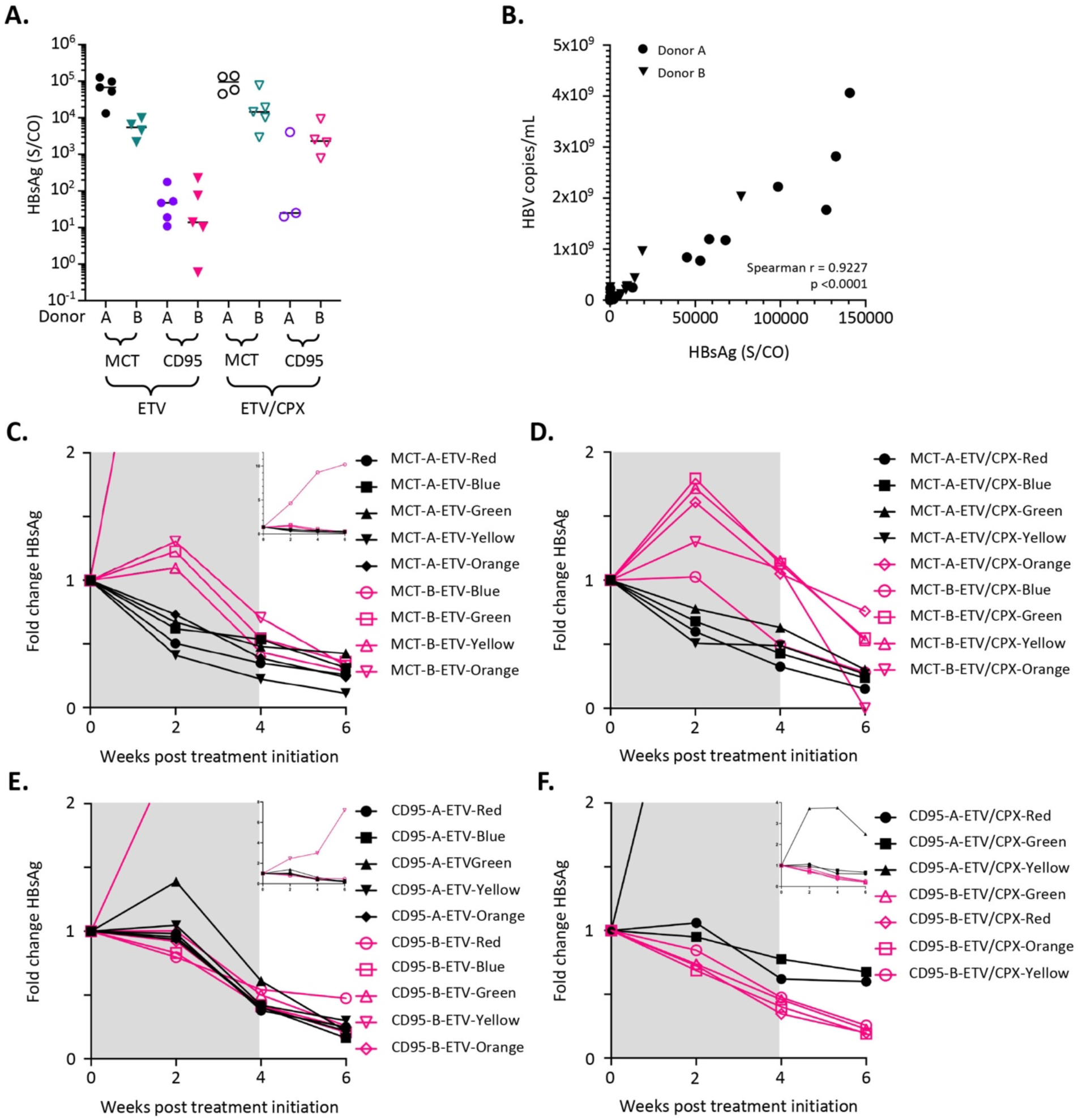
HBsAg levels in HBV infected liver humanized NSG-PiZ mice. At 10 weeks post-transplant humanized NSG-PiZ mice preconditioned with MCT or anti-CD95 mAb and transplanted with PHH from donor A or donor B, were challenged with 10^6^ copies of a Genotype D clinical isolate. Peak serum HBsAg levels were measured at week 10 post infection prior to antiviral administration (**A**). Spearman correlation between serum HBsAg levels and viral loads at 10 weeks post infection in 35 mice receiving PHH from donors A and B (**B**). Fold change in HBsAg levels were determined after antiviral treatment in individual mice pre-treated with MCT (**C**/**D**) or anti-CD95 antibody (**E**/**F**). At 10 weeks post-infection, infected mice were administered antivirals for 4 weeks (Grey bars) and HBsAg fold changes were monitored for a total of 6 weeks. Mice were given either oral ETV alone (0.5mg/Kg) in drinking water for 4 weeks (**C**/**E**) or oral ETV plus daily IP injections of CPX (5mg/Kg) for 4 weeks (**D**/**F**). Outlier mice with atypical HBsAg values are shown (inset panel **C**/**E**/**F**).

### Quantification of intrahepatic total HBV DNA and cccDNA

At 24 weeks post HBV inoculation, mice were euthanized and levels of human RPP30 (hRPP30), mouse RPP30 (mRPP30), total HBV DNA and cccDNA in liver were quantified by droplet digital (dd)PCR or qPCR (total HBV DNA) so that levels of hepatocyte chimerism could be determined along with levels of PHH-associated total HBV DNA and cccDNA. At necropsy, levels of human DNA (0.03-13.3%) and hepatocyte chimerism (0.08-33%) were variable, and 3 of 34 mouse livers had no detectable hRPP30 (**Figure 5A/B**). These 3 mice were all pre-treated with anti-CD95 antibody and had the lowest viral loads in serum at 10 weeks post HBV administration (3.6×10^4^-1.18×10^5^ HBV copies/ml). HBV was not detected in serum from these mice upon cessation of antiviral treatment until 4 weeks after antiviral treatment withdrawal. Levels of human DNA and PHH were higher in MCT-pretreated mice than in CD95 pretreated mice (**Figure 5A/B**), and all mice with detectable hRPP30 had detectable HBV DNA (0.04-84 copies/PHH, **Figure 5A/D**). Only 11 mice had detectable cccDNA (0.014-3.15 copies/PHH, **Figure 5E**), and all had total HBV DNA levels above 1.6 copies/PHH (11 of 15 mice with total HBV DNA levels above 1.6). At the time of death, a weak positive correlation was seen between levels of PHH and total HBV DNA (**Figure 5C**, Spearman r = 0.4657), but a strong positive correlation was seen between levels of total HBV DNA and levels of cccDNA **(Figure 5F**, Spearman r = 0.7165).

**Figure 5.**
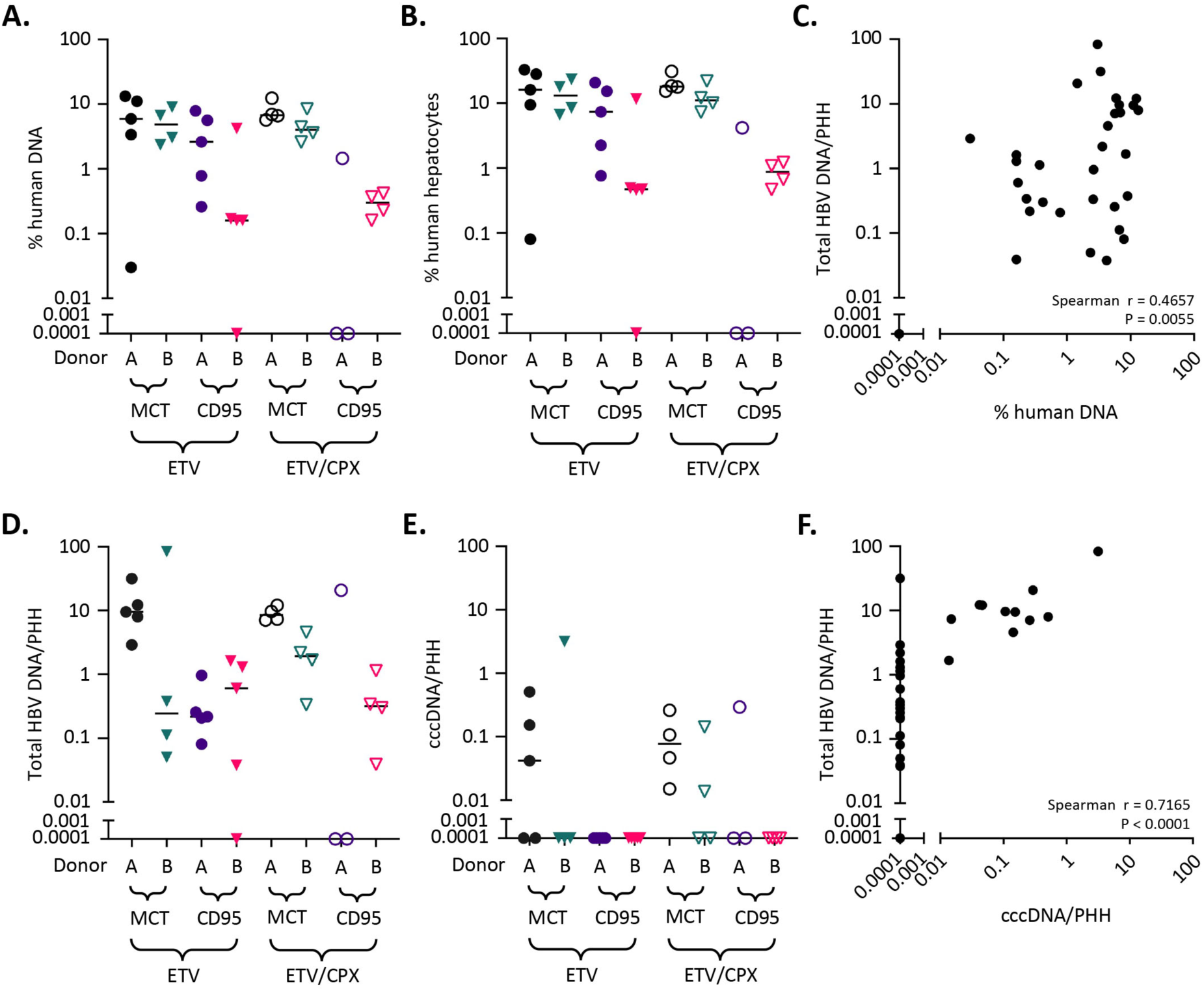
Human cell chimerism and liver-associated HBV DNA. Levels of cell-associated human RPP30, mouse RPP30, total HBV DNA and cccDNA in liver at necropsy (24 weeks post HBV) were quantified by ddPCR or qPCR (Total HBV DNA) using DNA extracted via Qiagen kit or modified Hirt extraction (cccDNA). Levels of human DNA (**A**), human hepatocyte chimerism (**B**), total HBV DNA (**D**) and cccDNA (**E**) were determined for all livers. Spearman correlation coefficients were calculated to determine whether levels of total cell-associated HBV DNA correlated with levels of human DNA (**C**) or cccDNA (F).

### AAV transduction of HBV+ PHH in liver humanized NSG-PiZ mice

Adeno-associated virus (AAV) vectors can deliver antiviral therapeutics to hepatocytes for the treatment of HBV ^36, 37^. To determine whether HBV+ PHH in our model can be transduced by AAV vectors, we administered scAAV.LK03-smCBA-GFP or scAAV3B-smCBA-GFP, which have known tropism for PHH in liver-humanized FRG and NSG-PiZ mice respectively ^56–58^, intravenously at doses of 1×10^12^, 2×10^11^ and 2×10^10^ vg/mouse to humanized NSG-PiZ mice that had been chronically infected with HBV for 142 days (n=3 per dose and serotype). PHH-naïve NSG-PiZ mice were only administered the highest dose for each AAV vector (n = 1 per serotype). GFP positive mouse hepatocytes were seen throughout livers of PHH-naïve NSG-PiZ mice 4 weeks after administration of scAAV.LK03-smCBA-GFP or scAAV3B-smCBA-GFP (**Figure 6A**). In HBV+ mice receiving 10^12^ vg scAAV.LK03-smCBA-GFP per animal, GFP positive mouse hepatocytes were also seen throughout the liver, but GFP expression was highly enriched in hCK18+/HBsAg+ PHH (**Figure 6B**). In contrast, scAAV3B-smCBA-GFP treated PHH+/HBV+ NSG-PiZ mice had GFP-positive mouse hepatocytes throughout the liver at 1×10^12^ vg/mouse, but GFP expression was not enriched in PHH, which contrasts a previous report of enriched PHH transduction by an AAV3B vector in humanized NSG-PiZ mice ^58^. GFP+/hCK18+/HBsAg+ PHH were detected, but at much lower frequency than in mice receiving AAV.LK03-smCBA-GFP (**Figure 6C**). For both AAV serotypes, fewer mouse hepatocytes and PHH were transduced at dose of 2×10^11^ and 2×10^10^ vg/mouse (data not shown).

**Figure 6.**
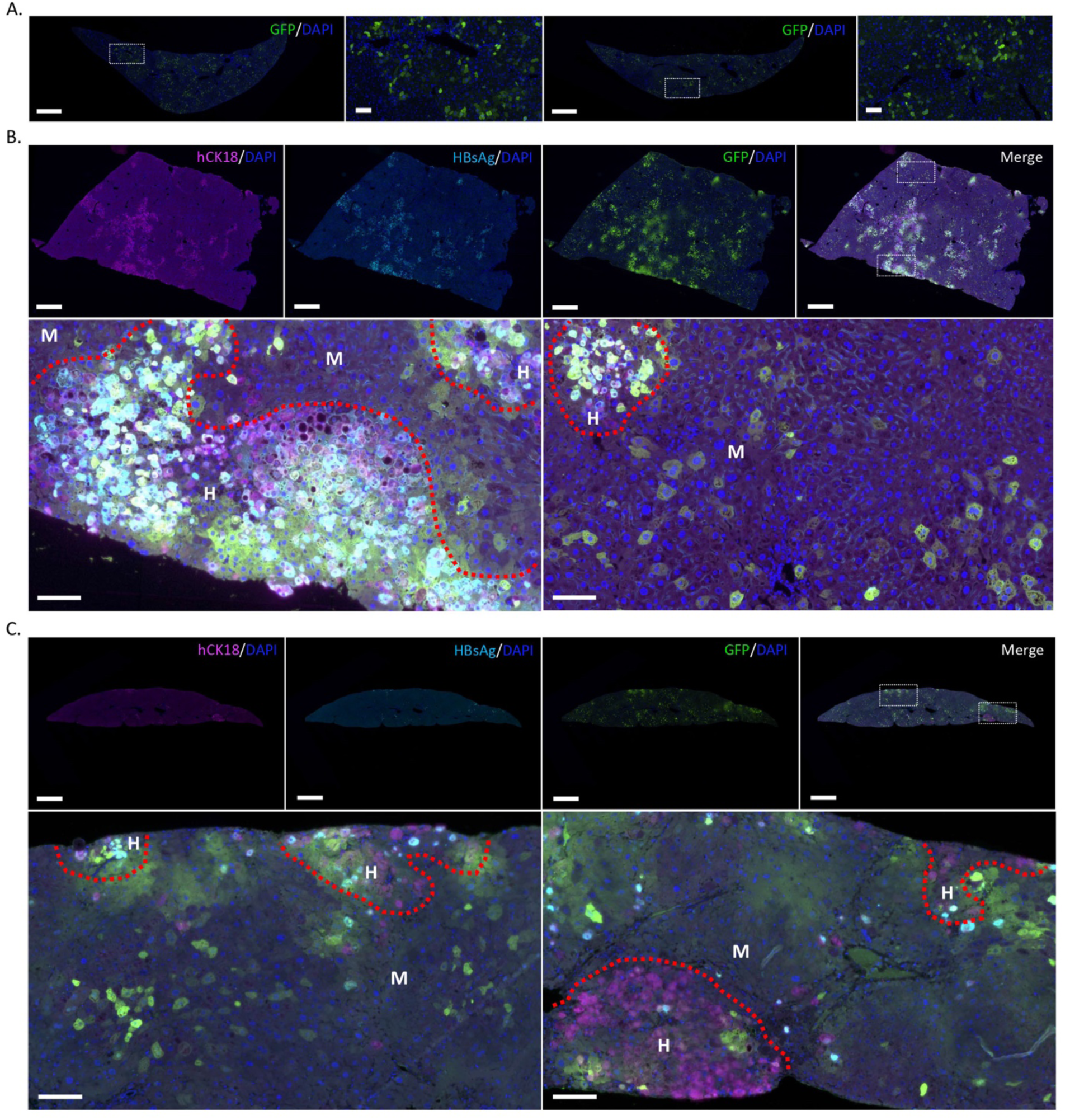
AAV transduction of HBV+ PHH in liver humanized NSG-PiZ mice. (**A**) Immunofluorescence imaging of GFP expression in control livers from PHH- NSG-PiZ mice 28 days after retro-orbital administration of 10^12^ vg of scAAV.LK03-smCBA-GFP (left hand images) or scAAV3b- smCBA-GFP. Scale bars 1mm (left image) or 100μm (right image). (**B**/**C**) Immunofluorescence labelling of GFP (green), hCK18 (purple) and HBsAg (cyan) in livers from NSG-PiZ mice pretreated with MCT, transplanted with 5×10^5^ primary human hepatocytes, challenged with 10^6^ IU of HBV, then administered 10^12^ vg of scAAV.LK03-smCBA-GFP (**B**) or scAAV3b-smCBA-GFP (**C**). One mouse had hAlb levels of 1.01 mg/mL at 12 weeks post PHH transplant and a viral load of 2.53 ×10^8^ IU/mL at 10 weeks post HBV challenge (**B**) and the other mouse had hAlb levels of 0.8 mg/mL at 12 weeks post PHH transplant and a viral load of 7.74 ×10^8^ IU/mL at 10 weeks post HBV challenge (**C**). For both mice, AAV was administered at 142 days post HBV challenge (213 days post PHH transplant) and euthanasia was performed at 27 days post AAV administration (169 days post HBV challenge, 240 days post PHH transplant). Areas of human (H) and mouse (M) hepatocytes are delineated by a red dotted line. Scale bars 1mm (upper images) or 100μm (lower images).

## Discussion

In this manuscript, we demonstrate that liver humanized NSG-PiZ mice are fully permissive to infection by HBV which selectively replicates in engrafted human hepatocytes and can establish chronic infections lasting at least 6 months. Infected livers contain cccDNA, secrete HBsAg and produce infectious virus that has now been serially passaged through three generations of mice without loss of infectivity. We also show that viral loads in chronically infected mice are reduced following treatment with the RTi entecavir, a front-line therapy for patients with CHB, and that HBV infected PHH can be efficiently transduced by AAV vectors, which allows this model to be used in the study of novel small molecule antivirals or gene-based curative therapies targeting HBV.

Liver humanized mice have become the gold standard for the study of HBV replication and therapy, and several liver humanized mouse models are available ^6, 15^. However, the complexity and/or cost of the most widely used uPA-SCID, FRG and TK-NOG models has limited their use and alternative models are still needed. We therefore investigated the NSG-PiZ mouse since it has several attributes that suggest it could be a robust and alternative model for the study of HBV. NSG-PiZ mice can be easily humanized via intra-splenic PHH injection ^34, 35, 58^, are readily available and affordable from commercial vendors, are viable and fertile, do not require a breeding license for academic use, are homozygous for the *SERPINA1* PiZ allele ensuring all-transgenic offspring, do not die as neonates without PHH transplantation due to toxic transgene expression, and do not require adenovirus-uPA administration or drug conditioning post-transplant to promote high level PHH engraftment. We now demonstrate that liver humanized NSG-PiZ mice are fully permissive to HBV infection, which can persist for at least 6 months. Levels of HBV replication are comparable to other liver humanized mouse models and strongly correlate with levels of PHH engraftment, and active infections can be established in mice with relatively low levels of PHH engraftment (huAlb > 0.04 mg/mL or <1% PHH engraftment). Thus, NSG-PiZ mice may be used to study disease progression and antiviral therapies during both acute and chronic HBV infections.

During our studies, we noticed that the kinetics of HBV replication were delayed in mice receiving PHH from donor B relative to donor A, although the levels of peak viremia achieved were similar. Delayed replication was more obvious in mice transplanted with fewer PHH (5×10^5^ vs 10^6^) and occurred despite donor B mice having ∼2-fold higher huAlb levels, a similar decrease in huAlb levels over 30 weeks of monitoring, and equivalent levels of human DNA at necropsy relative to donor A mice. This suggests that donor B PHH are, at least initially, less permissive to HBV infection, which is supported by our observations in HBV+ livers at necropsy. In donor A transplanted mice, the majority of hCK18+ PHH were HBsAg+ with a diffuse staining pattern observed in areas of engrafted PHH, whereas donor B transplanted mice contained HBsAg+ replication foci that appeared to be spreading through areas of uninfected hCK18+ PHH. The majority of HBsAg+ PHH within these foci were either undergoing macrovesicular steatosis or were adjacent to HBsAg+ PHH that were undergoing macrovesicular steatosis, which presents a potentially novel mechanism of viral spread for HBV. It has previously been shown that polyploid and binuclear hepatocytes are more prevalent in patients with more severe CHB infections^59^ and it is possible that HBV spread due to hepatocyte fusion is occurring in mice containing donor B PHH that are undergoing macrovesicular steatosis. Whether the observed viral spread in mice receiving donor B PHH is receptor-dependent or independent is yet to be determined and future studies will establish the role played by NTCP or other factors including host restriction factors that limit HBV replication.

Overall, our results provide broad evidence that liver humanized NSG-PiZ mice offer a robust alternative to existing models for the study of CHB and novel curative therapies. Furthermore, this model could enable broader access to HBV research in a more clinically relevant mouse model of disease.

## Acknowledgements

We thank Jennifer Duncan for helping establish the humanized NSG-PiZ model. We thank Amanda Koehne and Christine Watson for help with tissue pathology analysis.

## Contributions

RCT, DS, MA and KRJ conceived and designed the studies. RCT, DS, MAL, LK, PR, TKS, HX, LS, SLU, GP, MLH conducted experiments and acquired and analyzed data. RCT, DS and KRJ wrote the paper.

## List of abbreviations

AAV: adeno-associated virus
cccDNA: covalently closed circular DNA
CHB: chronic hepatitis B
CPX: ciclopirox
ddPCR: droplet digital PCR
ETV: entecavir
GFP: green fluorescent protein
HBcAg: hepatitis B core antigen
HBeAg: hepatitis B e antigen
HBsAg: hepatitis B surface antigen
HBV: hepatitis B virus
HCC: hepatocellular carcinoma
hRPP30: human RPP30
hCK18: human cytokeratin 18
HEK: human embryonic kidney
HHV: human herpesvirus
HPV: human papilloma virus
HSC: hematopoietic stem cells
HSV: herpes simplex virus
huAlb: human albumin
IHC: immunohistochemistry
MCT: monocrotaline
mRPP30: mouse RPP30
NSG: NOD-scid-gamma
NTCP: sodium taurocholate co-transporting polypeptide
PBS: phosphate buffered saline
PHH: primary human hepatocytes
qPCR: quantitative PCR
rcDNA: relaxed circular DNA
RTi: reverse transcriptase inhibitor
uPA: urokinase plasminogen activator

---

**Supplemental Figure 1.**
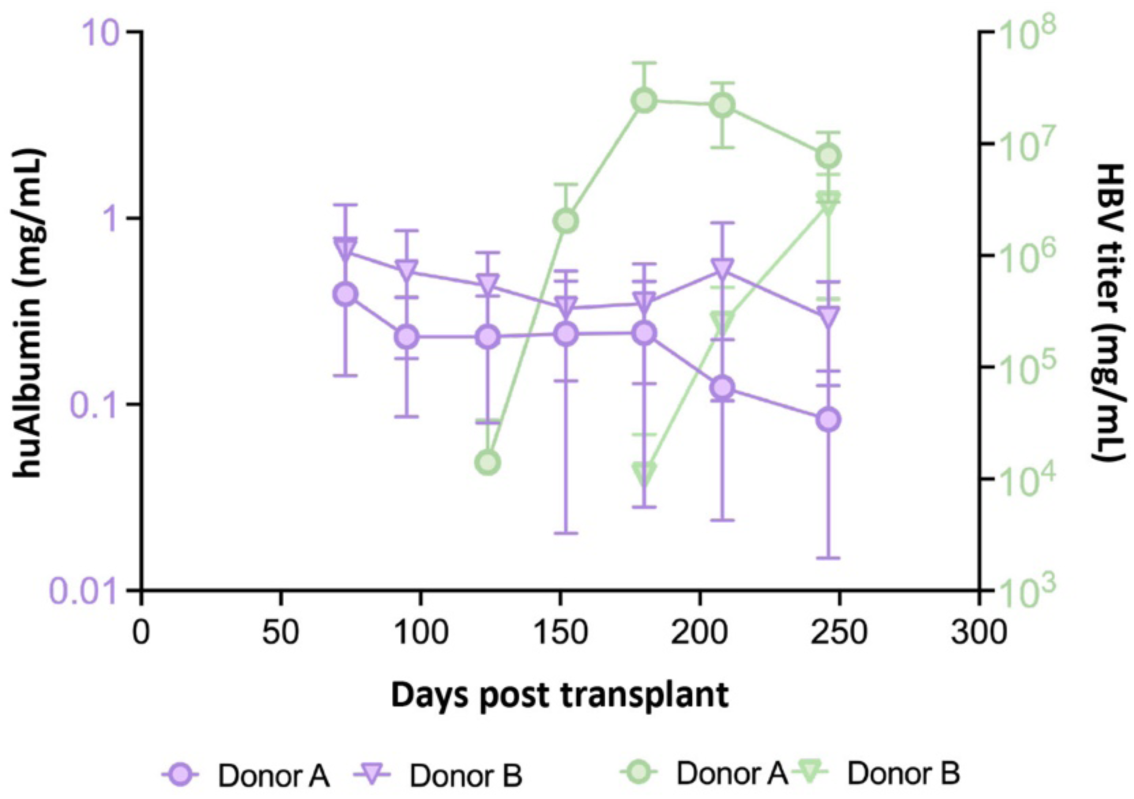
Longitudinal PHH engraftment and HBV replication in a pilot study performed in liver humanized NSG-PiZ mice. Mean serum huAlb levels (purple) and viral loads (green) following intravenous challenge with 10^6^ infectious units of a genotype D clinical isolate.

**Supplemental Figure 2.**
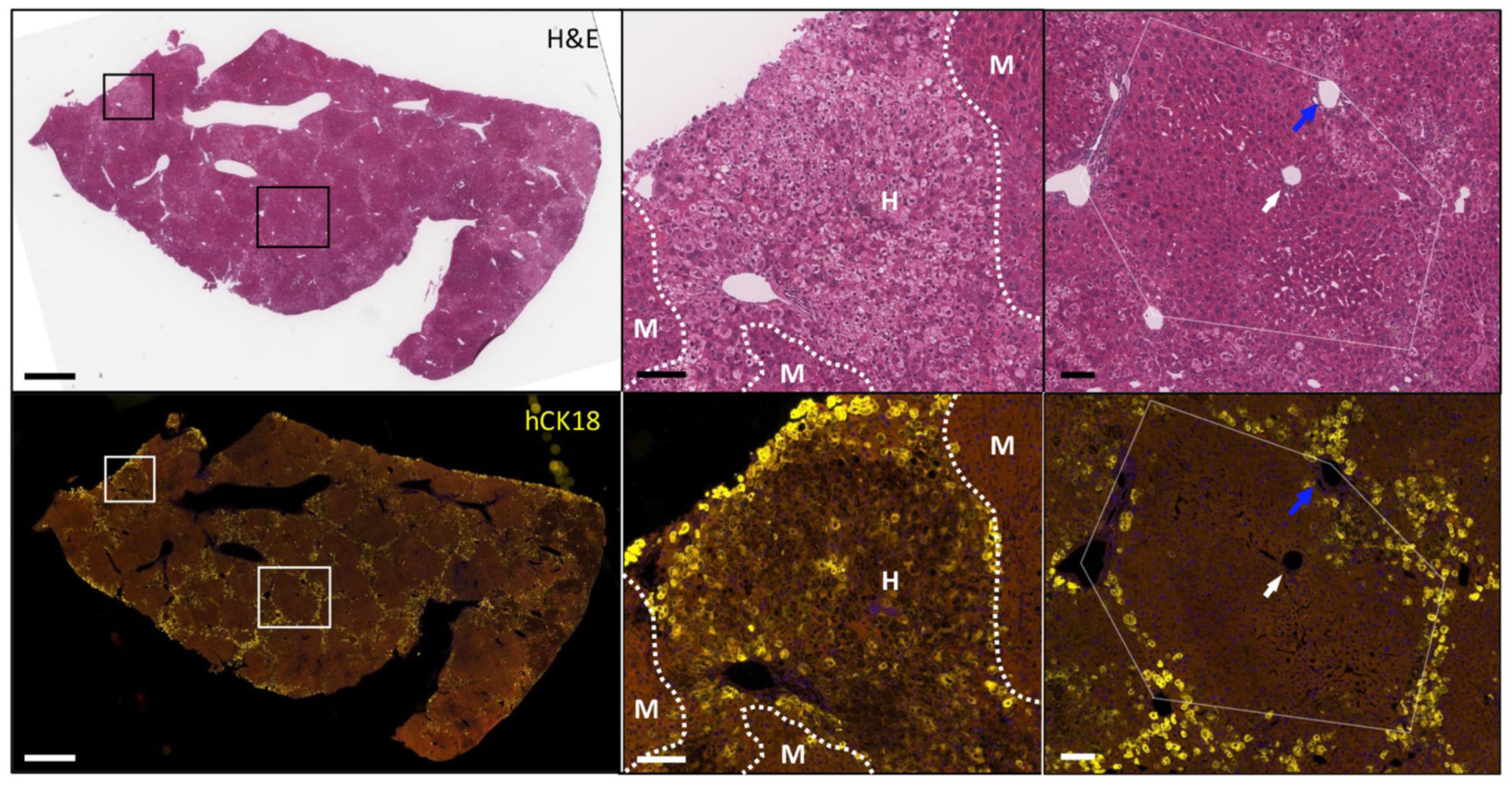
Human hepatocyte engraftment in NSG-PiZ mouse livers. H&E staining (Upper images) and hCK18 immunofluorescence (lower images) in serial sections from the left lateral lobe of an NSG-PiZ mouse pretreated with MCT, transplanted with 5×10^5^ primary human hepatocytes, and euthanized at 231 days post-transplant. This mouse had hAlb levels of 4.21 mg/mL at 8 weeks post hepatocyte transplant. Areas of liver containing human (H) or mouse (M) hepatocytes are shown (central images). At necropsy, human hepatocytes were predominantly periportal or surrounding hepatic lobules (right hand images), with few present within hepatic lobules or adjacent to the central vein. Scale bars 1mm (left hand images), 100μm (central and right-hand images).

**Supplemental Fig 3.**
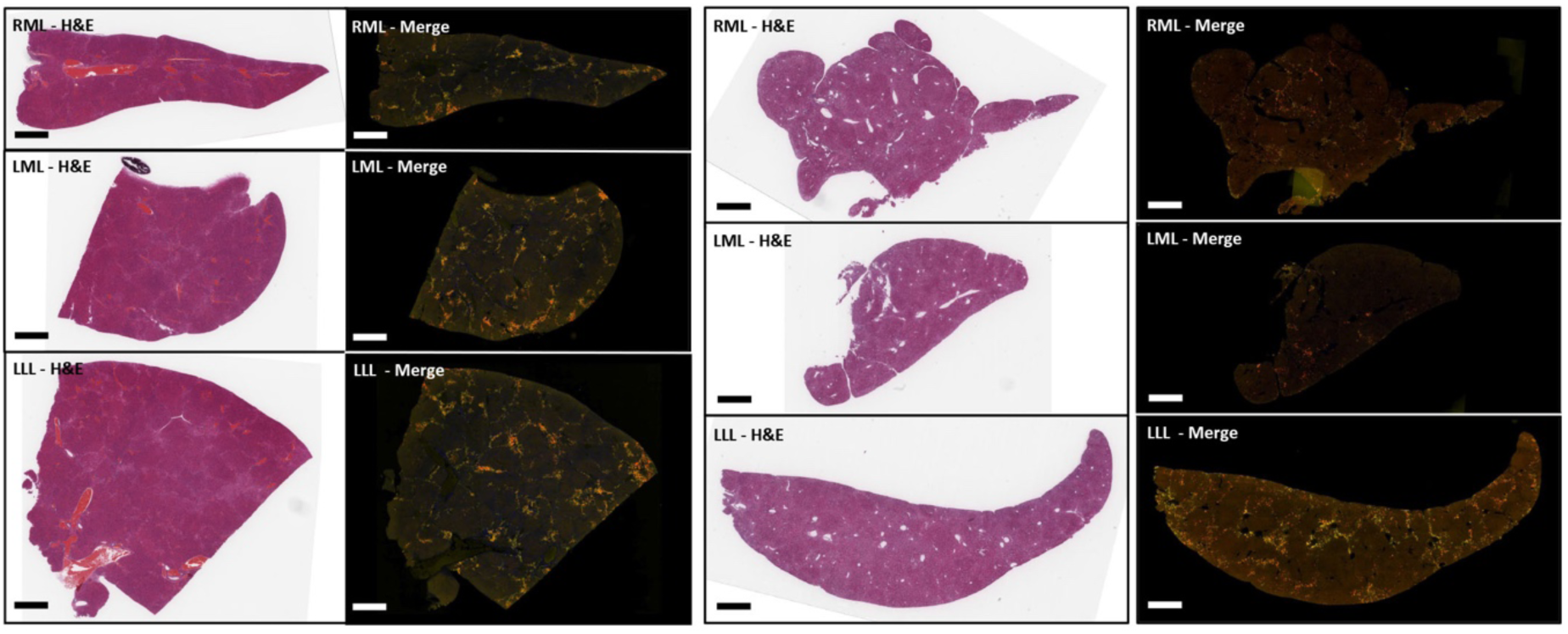
HBV replication occurs in all lobes of humanized NSG-PiZ mouse livers. The right medial (RML), left medial (LML) and left lateral (LLL) liver lobes are shown from two NSG-PiZ mouse pretreated with MCT, transplanted with 5×10^5^ PHH, and challenged intravenously with 10^6^ IU of a genotype D HBV clinical isolate. HBV challenge occurred at 10 weeks post PHH transplant and liver from a mouse that died at 150 days post HBV challenge is shown (upper images) alongside a mouse that was euthanized at 160 days post HBV challenge (lower images). Serial sections were stained with H&E or via immunofluorescence with antibodies against hCK18 (yellow) and HBsAg (Red). Scale bar - 1mm (left hand images) or 100μm (right hand images).

**Supplemental Figure 4.**
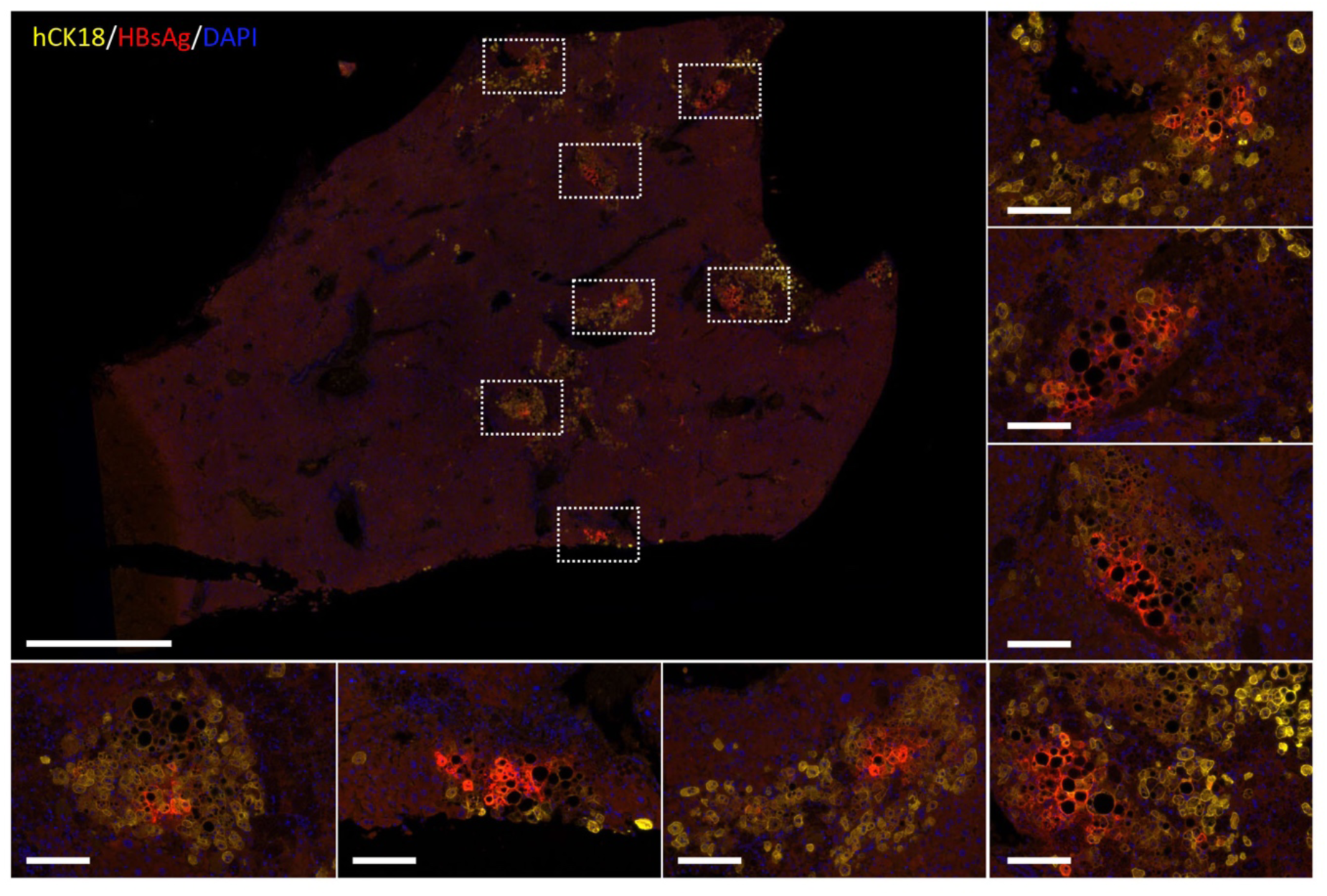
HBV replication in humanized NSG-PiZ mouse livers. HBsAg (red) and hCK18 (yellow) immunofluorescence in liver of an NSG-PiZ mouse pretreated with monocrotaline and transplanted with 5×10^5^ PHH from donor B, that died at 143 days post-transplant and 75 days post HBV challenge. Scale bars 1mm (large image), 100μm (smaller images).

**Supplemental Figure 5.**
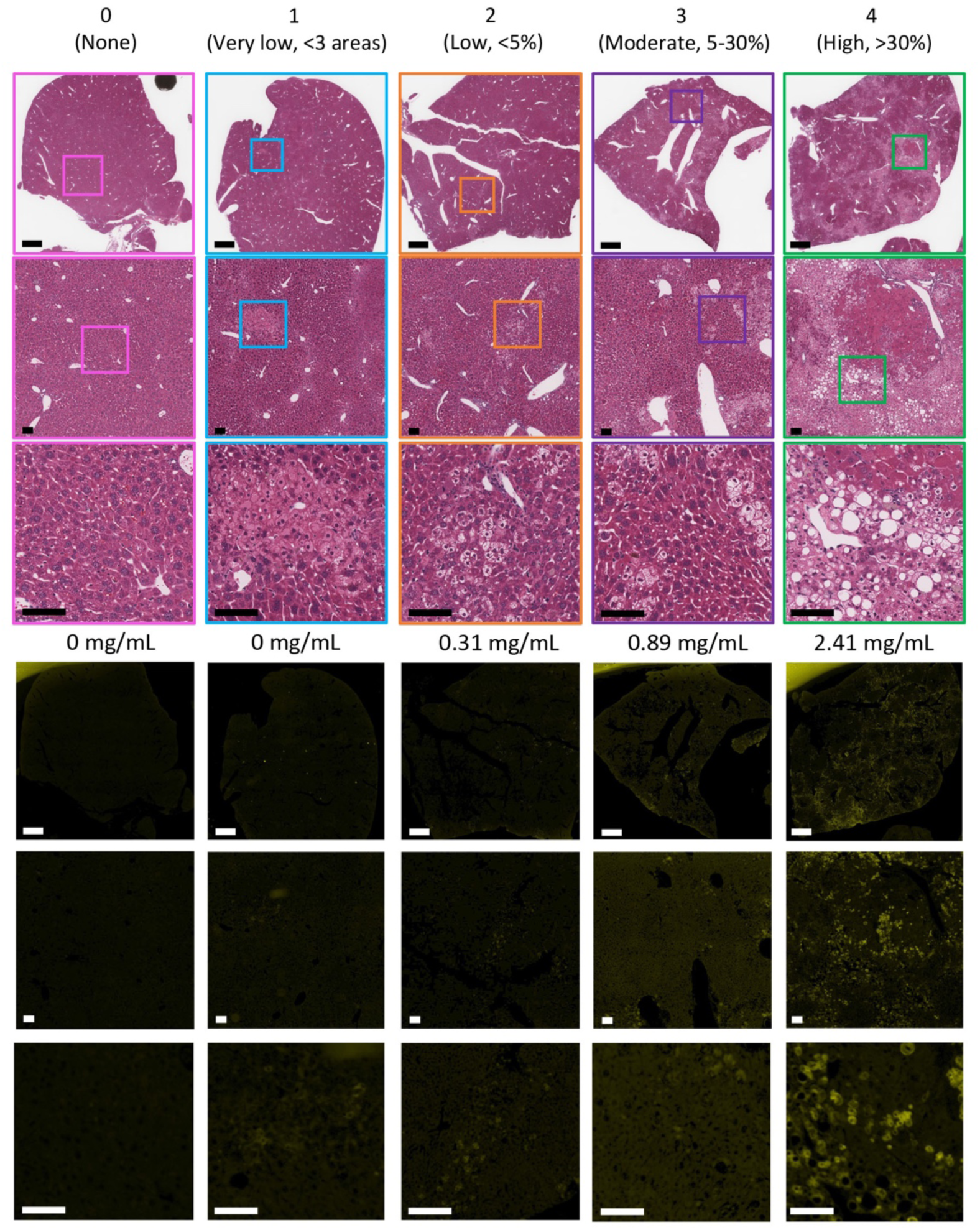
PHH NSG-PiZ engraftment scoring at necropsy for the experiment described in. **Figure 1.** Paraffin embedded sections from the left lateral, left medial and right medial liver lobes of each mouse were stained with H&E (Upper images) or for hCK18 expression by immunofluorescence (lower images). Each mouse was scored from 0 to 5 for levels of PHH engraftment at necropsy using H&E staining and levels of hCK18 staining as indicated. All mice were euthanized at day 242 post PHH transplantation except for 2 mice that died at days 183 and 215 post PHH transplant. The respective hAlbumin levels at 8 weeks post PHH transplant are shown for each example mouse. Scale bars 1mm (upper image) or 100μm (middle and lower images).

**Supplemental Figure 6.**
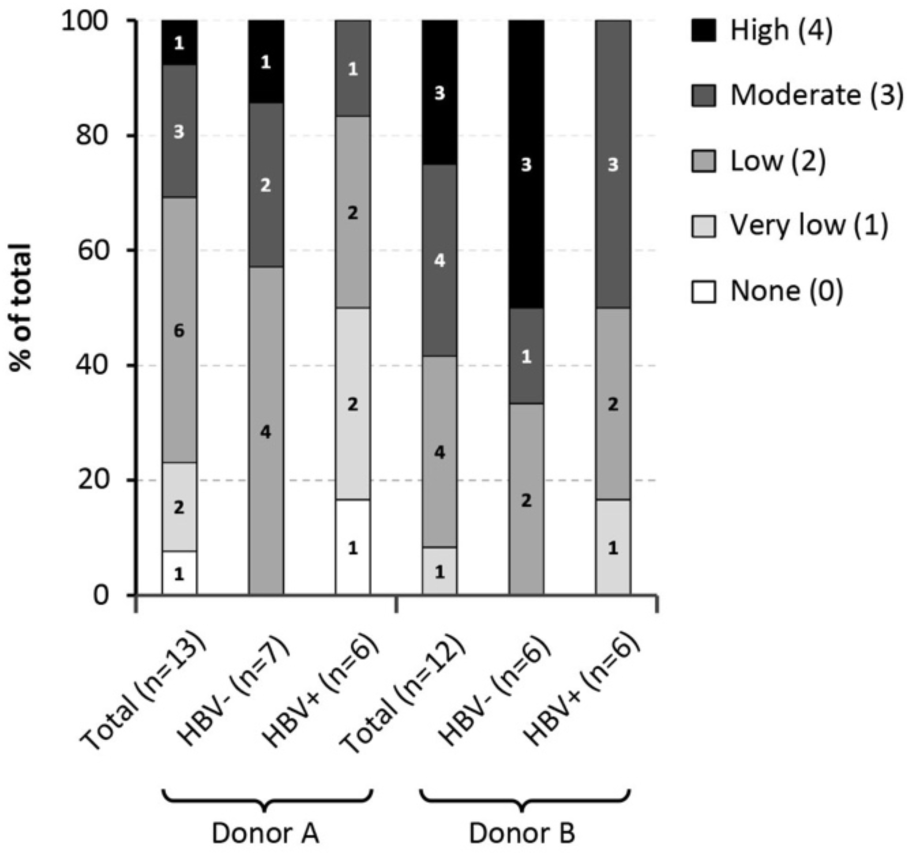
PHH NSG-PiZ engraftment scores at necropsy. All 25 mice from the experiment described in Figure 1 were scored from 0 to 5 for levels of PHH engraftment at necropsy using H&E staining and levels of hCK18 staining as indicated in Figure S5. All mice were euthanized at day 242 post PHH transplantation except for 2 mice that died at days 183 and 215 post PHH transplant.

**Supplemental Figure 7.**
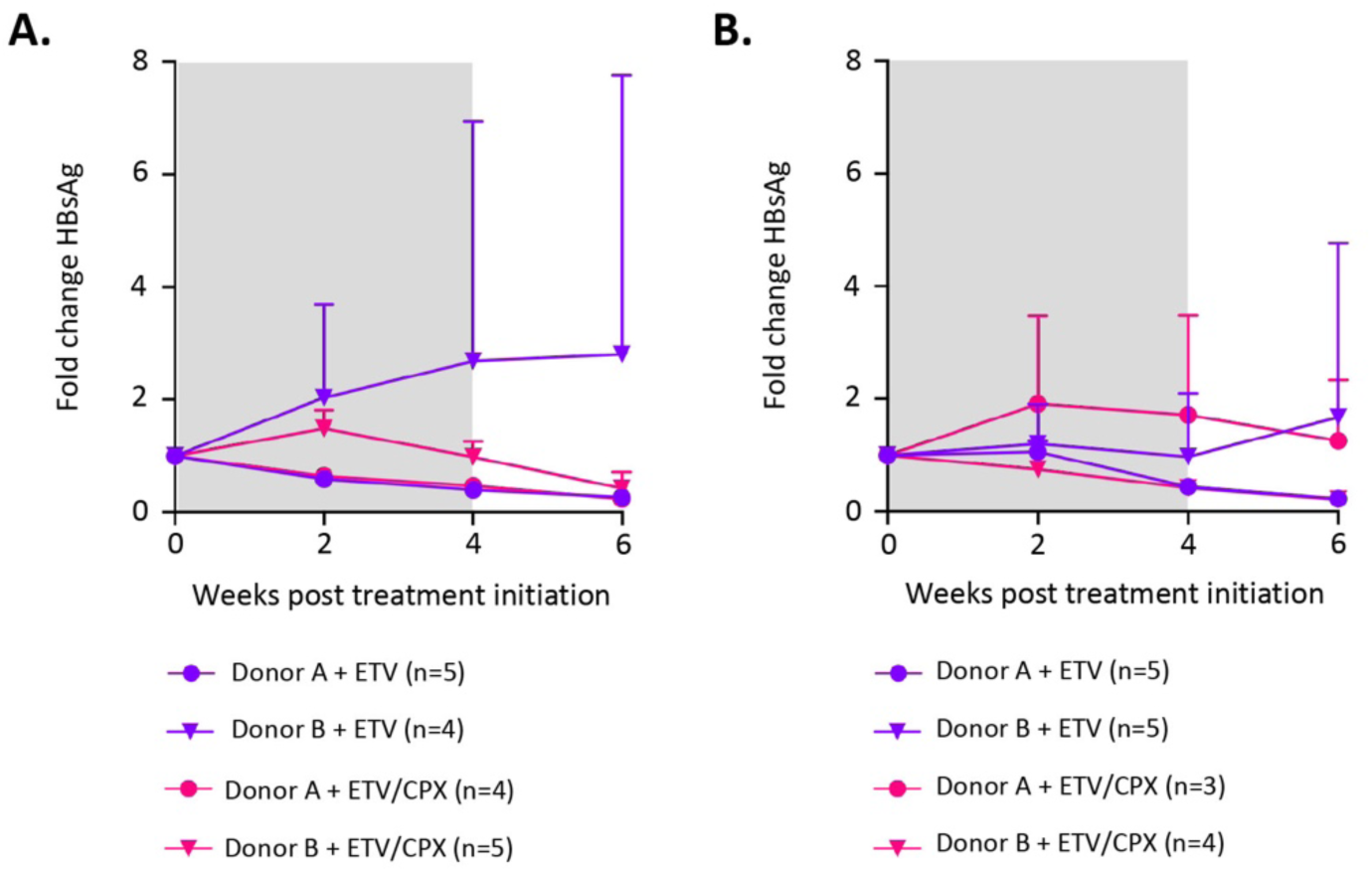
Mean fold change in HBsAg levels in HBV infected liver humanized NSG-PiZ mice after antiviral treatment. Serum HBsAg levels peaked at week 10 post infection and the fold change in HBsAg levels were determined for mice pre-treated with MCT (**A**) or anti-CD95 antibody (**B**) after antiviral treatment. At 10 weeks post-engraftment humanized NSG-PiZ mice were challenged with 10^6^ copies of a Genotype D clinical isolate. At 10 weeks post-infection, chronically infected mice were given ETV in drinking water (0.5mg/Kg) +/- daily IP injections of CPX (5mg/Kg) as indicated for 4 weeks (Grey bars). Error bars = SD.

